# Strain-dependent contribution of the AcrAB-TolC efflux pump to *Klebsiella pneumoniae* physiology

**DOI:** 10.1101/2025.09.26.678718

**Authors:** Kirandeep Bhogal, Barbara Clough, Charlotte Emmerson, Archie Organ, Yin Chen, Michelle MC Buckner, Ilyas Alav

**Affiliations:** Institute of Microbiology and Infection, University of Birmingham, Birmingham, UK; Department of Microbes, Infection and Microbiomes, School of Infection, Inflammation and Immunology, College of Medicine and Health, University of Birmingham, Birmingham, UK; School of Biosciences, College of Life and Environmental Sciences, University of Birmingham, Birmingham, UK

## Abstract

*Klebsiella pneumoniae* is a prominent opportunistic pathogen increasingly associated with multidrug resistance and virulence. One of the main mechanisms of antimicrobial resistance in *K. pneumoniae* is active efflux, primarily mediated by the Resistance-Nodulation-Division (RND) family of pumps. AcrAB-TolC is the key RND efflux pump in *K. pneumoniae*, regulated by the transcriptional activator RamA and its repressor RamR. Although overexpression of AcrAB-TolC has been linked to drug resistance in various clinical strains, its physiological roles in *K. pneumoniae* remain insufficiently studied. In this study, we generated isogenic deletions of *acrB* and *ramR* in both the genetically tractable *K. pneumoniae* Ecl8 and the virulent ATCC 43816 strains. We examined the phenotype of the Δ*acrB* and Δ*ramR* mutants by assessing antimicrobial susceptibility, biofilm formation, growth under infection-related conditions, and both *in vitro* and *in vivo* infection models. Loss of *acrB* increased susceptibility to drugs, decreased biofilm formation, and reduced *in vitro* virulence in both Ecl8 and ATCC 43816. However, only in Ecl8 was the loss of AcrB found to diminish growth under infection-like conditions and decrease *in vivo* virulence in the *Galleria mellonella* infection model. In contrast, in ATCC 43816, it had no effect. Our findings suggest that AcrAB-TolC exhibits strain-specific physiological functions, highlighting its dual role in antimicrobial resistance and pathogenicity, and thereby broadening our understanding of efflux-mediated adaptations in *K. pneumoniae*. Exploring the broader functions of RND efflux pumps in *K. pneumoniae* can provide insights into the potential effects of targeting them with inhibitor molecules.

**Importance:** Infections caused by multidrug-resistant bacterial pathogens are among the most urgent global public health challenges. Specifically, *Klebsiella pneumoniae* is classified by the World Health Organisation as a critical priority pathogen for the development of new treatments. Resistance-Nodulation-Division (RND) efflux pumps significantly contribute to multidrug resistance and virulence, making them promising targets for drug development. In this study, we demonstrated that the RND efflux pump AcrAB-TolC in *K. pneumoniae* is essential for drug efflux, biofilm formation, and *in vitro* virulence. Notably, we identified strain-specific roles for AcrAB-TolC in supporting growth under infection-related conditions and virulence in the *Galleria mellonella* infection model. Our findings highlight that the function of RND efflux pumps can vary between strains within a species. This implies that targeting RND efflux pumps with inhibitors may yield different effects depending on the strain background.

## Introduction

*Klebsiella pneumoniae* is a major cause of opportunistic and nosocomial infections worldwide (1). As a commensal member of the human microbiota, *K. pneumoniae* can exploit compromised host defences to cause various opportunistic infections, including urinary tract infections, pneumonia, wound infections, and bacteraemia (2). The occurrence and levels of antimicrobial resistance in *K. pneumoniae* have risen sharply over recent decades, resulting in its classification as a critical priority pathogen by the World Health Organisation (3, 4). Alarmingly, hypervirulent *K. pneumoniae* strains have emerged in recent years, exhibiting increased virulence and the ability to infect healthy individuals (5).

In *K. pneumoniae* and other Gram-negative pathogens, the resistance-nodulation-division (RND) superfamily efflux pumps span the entire cell envelope, conferring intrinsic resistance to a wide range of antimicrobial compounds (6). The archetypal RND efflux pump in Enterobacteriaceae is the AcrAB-TolC efflux system, comprised of the inner membrane transporter AcrB, the periplasmic adapter protein AcrA, and the outer membrane channel TolC (7). In clinical Gram-negative isolates, RND efflux pumps, such as AcrAB-TolC, are frequently overexpressed, thereby contributing to multidrug resistance by expelling antibiotics and reducing their intracellular accumulation (8). RamA is a key transcriptional activator of AcrAB-TolC expression in Enterobacteriaceae such as *K. pneumoniae* and *Salmonella enterica* (9, 10), with MarA as the equivalent in *Escherichia coli* (11). RamA activity is repressed by its local repressor RamR through binding to the *ramA* promoter region (12). RamR itself is regulated by environmental signals, including bile acids and antibiotics, which bind to RamR and decrease its affinity for DNA binding, including the *ramA* promoter (13). Therefore, inactivating mutations in *ramR* that render RamR non-functional result in significant RamA accumulation and AcrAB-TolC overexpression, thereby affecting the microbe-antibiotic response (14). For example, mutations in *ramR* have been identified in clinical *K. pneumoniae* isolates, contributing to tigecycline resistance by upregulating *acrAB*-*tolC* expression (15).

In addition to their role in multidrug resistance, RND efflux pumps, including AcrAB-TolC, play a role in bacterial virulence (16). In several species of Enterobacteriaceae, the loss of *acrB* confers reduced virulence *in vitro* and/or *in vivo* (17–19). In *K. pneumoniae* 52145R and A2312, deleting *acrB* has previously been shown to increase antimicrobial susceptibility and reduce virulence in mouse infection models (20, 21). However, the roles of the AcrAB-TolC efflux pump in *K. pneumoniae* Ecl8, a widely used reference strain for targeted genetic manipulation, and *K. pneumoniae* ATCC 43816, a hypervirulent reference strain widely used for studying pathogenesis, have not been studied (22, 23). Furthermore, the loss and overexpression of AcrAB-TolC activity in *K. pneumoniae* have not been studied in relation to biofilm formation, growth in infection-relevant conditions, and *in vitro* infection.

In this study, isogenic *acrB* and *ramR* deletion strains of *K. pneumoniae* Ecl8 and ATCC 43816 were generated. The phenotypes of the mutants were characterised by measuring antimicrobial susceptibility, biofilm formation, survival in human serum, growth in artificial lung media, and the ability to cause infection *in vitro* and the *in vivo Galleria mellonella* infection model. This study shows that whilst AcrAB-TolC plays a fundamental role in antimicrobial susceptibility in different *K. pneumoniae* strains, its impact on growth in infection-relevant conditions and in vivo virulence is strain-dependent. This could have potential implications for the targeting of AcrAB-TolC by inhibitors in different *K. pneumoniae* strains.

## Methods

### Bacterial strains

The bacterial strains used are described in Table 1. Unless stated otherwise, all strains were grown in Luria–Bertani (LB) broth (Sigma-Aldrich, USA) and incubated at 37 °C with aeration.

**Table 1.**
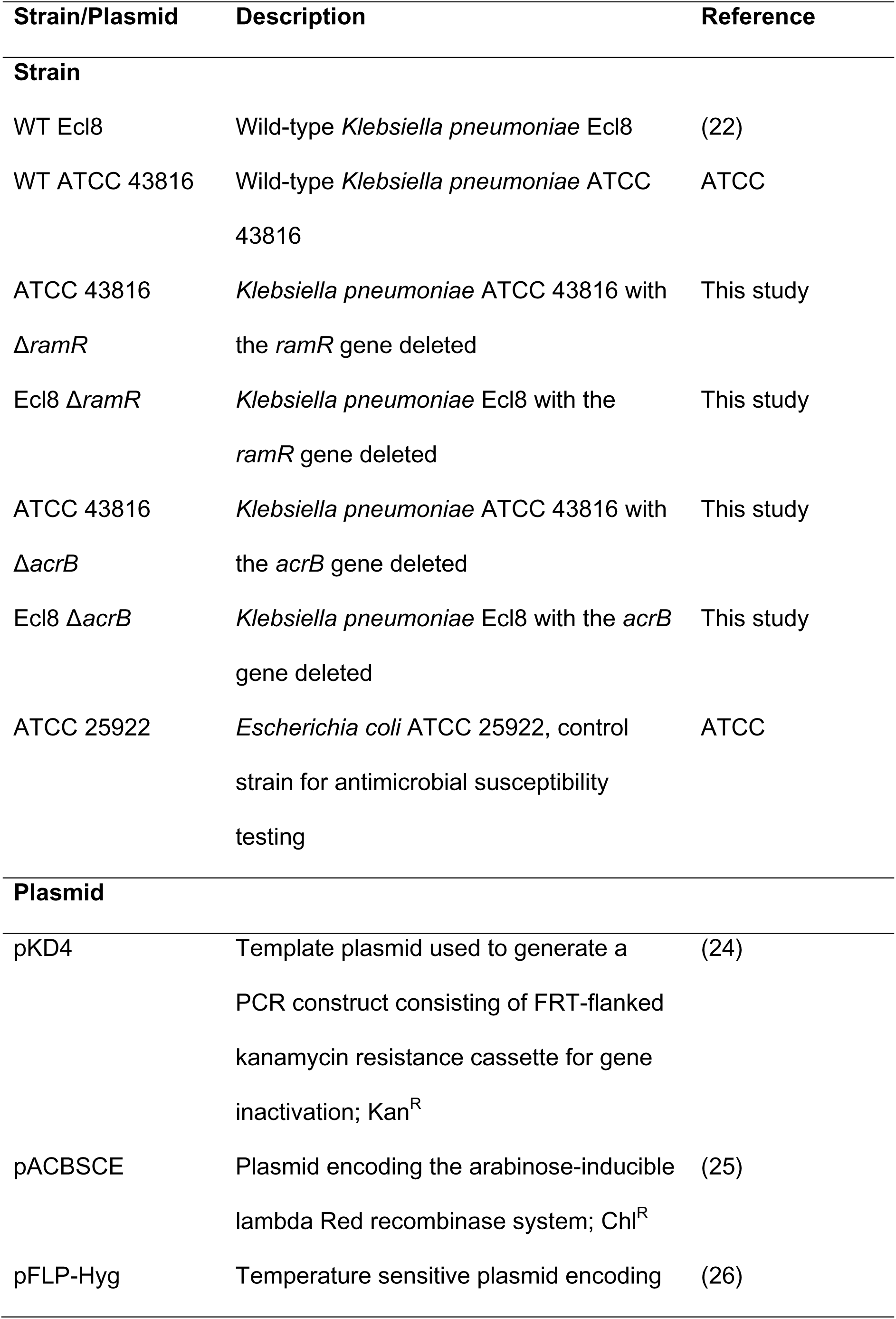
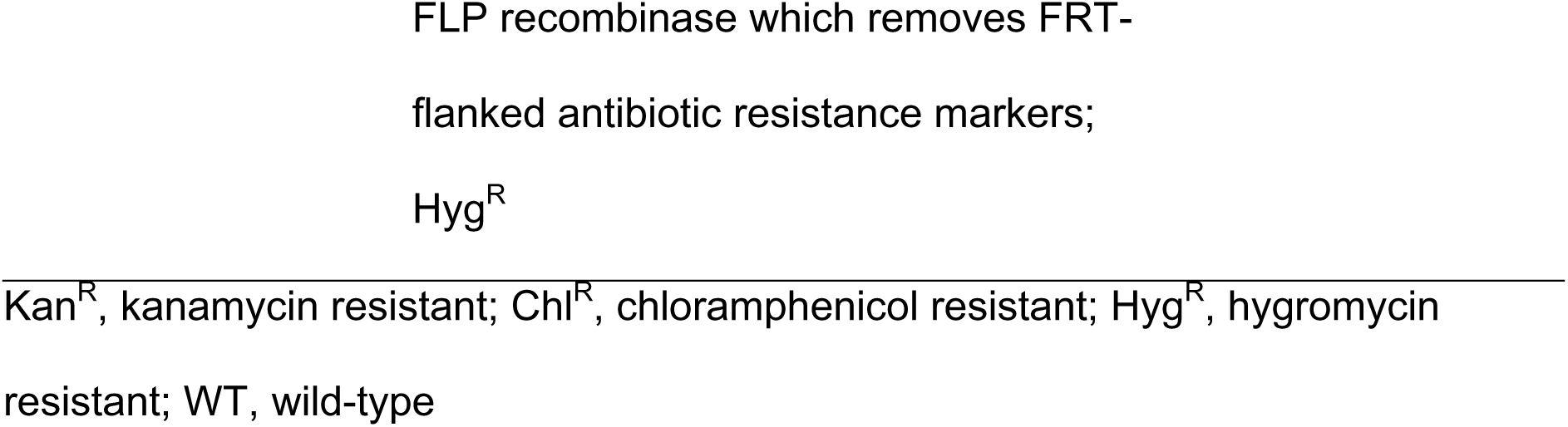
List of bacterial strains and plasmids used in this study.

### Generation of the *ramR* and *acrB* deletion mutant strains

The *acrB* and *ramR* mutant strains of Ecl8 and ATCC 43816 were constructed using λ Red recombination, followed by FLP recombinase to obtain antibiotic-susceptible gene knockouts (24). The *acrB* or *ramR* gene was inactivated by the insertion of the kanamycin resistance gene (*aph*) gene using the pACBSCE recombineering plasmid as described previously (25). The *aph* gene was subsequently removed using the pFLP-Hyg plasmid (gifted from Pep Charusanti; Addgene plasmid #87831; http://n2t.net/addgene:87831; RRID: Addgene_87831) as described previously (26). All primers used to generate the genetic deletions are listed in Supplementary Table 1.

### Relative RT-qPCR

All protocols were carried out according to the manufacturer’s instructions. Total RNA was extracted from *K. pneumoniae* Ecl8 and ATCC 43816 strains grown to the exponential phase (OD_600_ = 0.4–0.5) using the Monarch Total RNA Miniprep Kit (NEB, USA), with on-column DNase I treatment to eliminate genomic DNA contamination. RNA quality and quantity were determined using a NanoDrop spectrophometer (Thermo Scientific, USA) and the Qubit RNA Broad Range Assay Kit (Invitrogen, USA), respectively. cDNA was synthesised from 1 µg of total RNA using the iScript Reverse Transcription Supermix Kit (Bio-Rad, USA), with no-reverse transcriptase controls included for each sample to confirm the absence of genomic DNA contamination. Real-time PCR was performed using the SensiFAST SYBR Lo-Rox Kit (Meridian Bioscience, USA) with gene-specific primers (Supplementary Table 2) and 1 ng/µL of cDNA per reaction on a QuantStudio 1 Real-Time PCR System (Thermo Fisher Scientific, USA). The *rpoB* gene, encoding the β subunit of bacterial RNA polymerase, was used as an endogenous control for normalising gene expression using the 2^−ΔΔ*Ct*^ method (27). RT-qPCRs were carried out using three biological replicates per strain, each with three technical replicates.

### Antimicrobial susceptibility testing

The minimum inhibitory concentration of antibiotics and biocides was determined using the broth microdilution method according to Clinical and Laboratory Standards Institute guidelines (28). *Escherichia coli* ATCC 25922 was used as a control strain for antimicrobial susceptibility testing.

### Serum survival assays

Overnight cultures of *K. pneumoniae* Ecl8 and ATCC 43816 were sub-cultured (2% inoculum) in 5 mL LB broth and grown to mid-log phase (OD_600_ = 0.5; ∼5×10^8^ CFU/mL). Cultures were diluted to 1×10^6^ CFU/mL in PBS, and 20 µL of this suspension was mixed with 180 µL of normal human serum (Merck, USA) or heat-inactivated serum (56 °C, 30 min) in round-bottom, non-treated 96-well plates, yielding 1×10^5^ CFU/mL per well. Plates were incubated statically at 37 °C for 3 h, and viable bacteria were enumerated by serial dilution and plating. Each strain was tested in three biological replicates with three technical replicates each.

### Ethidium bromide efflux assays

The efflux activity of *K. pneumoniae* Ecl8 and ATCC 43816 strains was determined using the ethidium bromide efflux assay as described previously (29).

### Crystal violet biofilm assays

Biofilm formation by *K. pneumoniae* Ecl8 and ATCC 43816 strains was measured as previously described (30). For each strain, three biological replicates were tested, each consisting of three technical replicates, conducted on separate occasions.

### Growth kinetic assays

Overnight cultures of *K. pneumoniae* Ecl8 and ATCC 43816 strains were adjusted to an OD_600_ of 0.01 (∼1×10^7^ CFU/mL). A 180 µL volume of the growth media was added to the wells of a flat-bottom non-treated 96-well polystyrene plate (Corning, USA), and 20 µL of the OD_600_-adjusted bacterial cells were added to the wells. The OD_600_ was recorded at 30-minute intervals over 18 hours using a FLUOstar Optima plate reader (BMG Labtech, Germany). For each strain, three biological replicates were tested, each consisting of three technical replicates, conducted on separate occasions.

### RAW 264.7 macrophage infection assays

RAW 264.7 macrophages (ATCC TIB-71) were cultured in DMEM with GlutaMAX (Thermo Fisher, USA) supplemented with 10% heat-inactivated FBS (Life Technologies, USA) at 37 °C and 5% CO_2_. Cells were seeded at 1×10^5^ per well in 96-well flat-bottom plates and infected in triplicate with exponential-phase bacteria at a multiplicity of infection (MOI) of 10. Plates were centrifuged (900 × *g*, 5 min) to synchronise infection, and extracellular bacteria were killed after 30 min with 200 µg/mL gentamicin, which remained in the medium thereafter. Uptake was quantified at 30 min, 1 h, and 2 h, and survival at 24 h post-infection by washing cells in PBS, lysing with 0.1% Triton X-100, serially diluting, and plating on LB agar. CFUs were enumerated after overnight incubation at 37 °C. Each strain was tested with three biological replicates across three independent experiments.

### A549 lung epithelial cell infection assays

A549 lung epithelial cells were cultured in RPMI with GlutaMAX (Thermo Fisher, USA) and 10% heat-inactivated FBS at 37 °C and 5% CO_2_. Cells were seeded at 3×10^4^ per well in 96-well flat-bottom plates and infected the following day with exponential-phase bacteria at an MOI of 50 for 2 h. Subsequent processing was performed as described above. Each strain was tested with three biological replicates across three independent experiments.

### *Galleria mellonella* larvae infection model

*Galleria mellonella* larvae were purchased from Livefood UK and stored at 15 °C in darkness with a non-restricted diet. Larvae were injected (*n* = 10 per strain, which was independently repeated three times) with 5×10^5^ bacterial cells as previously described (31), and the number of live/dead larvae was quantified over 3 days.

### Growth in healthy lung media (HLM) and cystic fibrosis lung media (CFLM)

Artificial media mimicking healthy lung (HLM) and cystic fibrosis lung (CFLM) environments were prepared as described previously (32). Overnight LB cultures were pelleted, resuspended in PBS, and diluted to 5×10^7^ CFU/mL in HLM or CFLM. In 96-well flat-bottom plates, 20 µL of bacterial suspension was mixed with 180 µL of HLM or CFLM (final inoculum of 5×10^6^ CFU/mL) and incubated at 37 °C with 5% CO₂. At 2, 6, 24, and 48 h, 10 µL samples were serially diluted in PBS and plated on LB agar. Colonies were enumerated after overnight incubation at 37 °C. Each strain was tested with three biological replicates across three independent experiments.

## Results

### The effect of *acrB* deletion and overexpression on efflux activity and antimicrobial susceptibility of *Klebsiella pneumoniae* Ecl8 and ATCC 43816

The deletion of *acrB* or *ramR* in both strains did not affect growth in LB broth or cation-adjusted Mueller-Hinton broth (Fig. S1). RT-qPCR was used to verify that the loss of *ramR* resulted in overexpression of *ramA* and the *acrAB*-*tolC* genes. In both Ecl8 and ATCC 43816, the deletion of *ramR* significantly increased the expression of *acrA*, *acrB* and *tolC* (Fig. S2). As expected (10), the expression of *ramA* was significantly upregulated in both strains, with expression being substantially higher in the ATCC 43816 strain (Fig. S2). Active efflux of ethidium bromide was measured by pre-loading cells with the dye and tracking the decrease in fluorescence as the substrate was exported. Efflux capacity was determined by the time taken for fluorescence to decrease to 50% of its initial value. It took significantly longer for the *K. pneumoniae* Ecl8 Δ*acrB* and ATCC 43816 Δ*acrB* strains to reduce fluorescence by 50% compared to their respective wild-type strains (Fig. 1). This demonstrated that the Δ*acrB* mutant strains had impaired efflux function. The Δ*ramR* strains took significantly less time to reduce the amount of fluorescence by 50% of the starting value compared to their respective wild-type strain, indicating increased AcrAB-TolC activity (Fig. 1).

**Figure 1.**
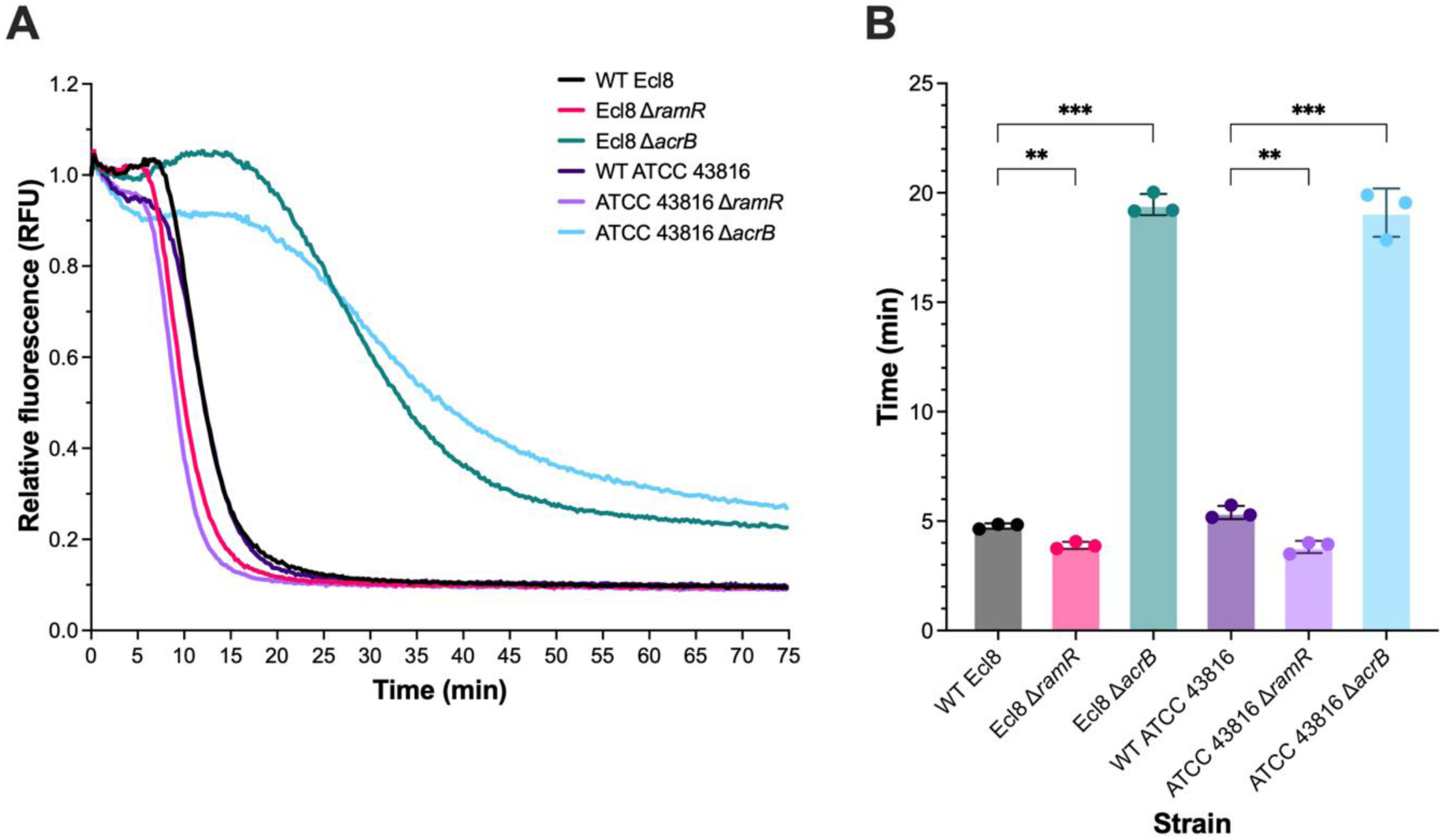
Efflux of ethidium bromide over time by *Klebsiella pneumoniae* Ecl8 and ATCC 43816 strains. Bacteria were treated with the efflux substrate ethidium bromide and the protonophore CCCP for 1 h, and then re-energised with glucose. **A)** Decrease in ethidium bromide fluorescence over time after re-energisation with glucose. The data shown represent the mean of three biological replicates, conducted on independent occasions. **B)** Time taken for ethidium bromide fluorescence to decrease by 50% of the starting value. The data shown represent the mean ± standard deviation of three biological replicates, conducted on independent occasions. Statistical significance was determined by comparing the wild-type Ecl8 or ATCC 43816 to their isogenic mutant strains using one-way ANOVA, followed by Dunnett’s test to correct for multiple comparisons. Significantly different results are indicated with ** (P ≤ 0.01) or *** (P ≤ 0.001).

The deletion of *acrB* in *K. pneumoniae* in Ecl8 and ATCC 43816 increased susceptibility to a broad range of well-known AcrB substrates, as indicated by a two-fold or greater reduction in MIC values compared to their respective wild-type strains. These included several different classes of antibiotics, such as quinolones (ciprofloxacin and nalidixic acid), chloramphenicol, macrolides (azithromycin and erythromycin), and tetracyclines (minocycline, tetracycline and tigecycline), biocides (benzalkonium chloride, chlorhexidine digluconate, and triclosan), and dyes (acriflavine, crystal violet and ethidium bromide). There was no difference in susceptibility between the Δ*acrB* and wild-type strains to several different β-lactams, including ceftazidime, cefepime, piperacillin, and meropenem. Like AcrB in *E. coli* and *S.* Typhimurium (33, 34), the loss of *acrB* in Ecl8 and ATCC 43816 did not change susceptibility to aminoglycosides, including amikacin and gentamicin (Table 2). The loss of *acrB* in *K. pneumoniae* Ecl8 and ATCC 43816 reduced MICs of several biocides, including benzalkonium chloride, chlorhexidine digluconate, and triclosan, but not octenidine.

**Table 2.**
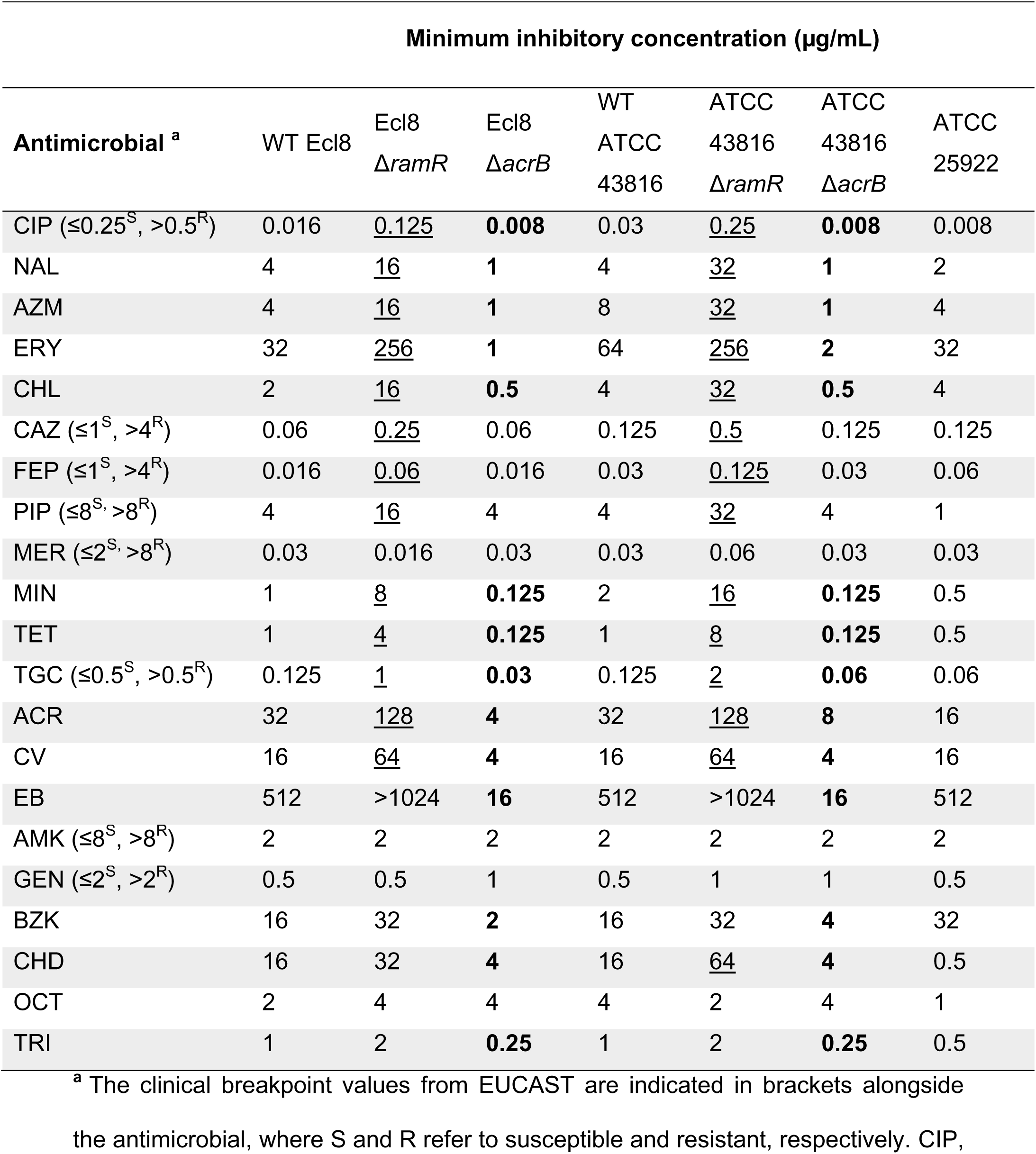
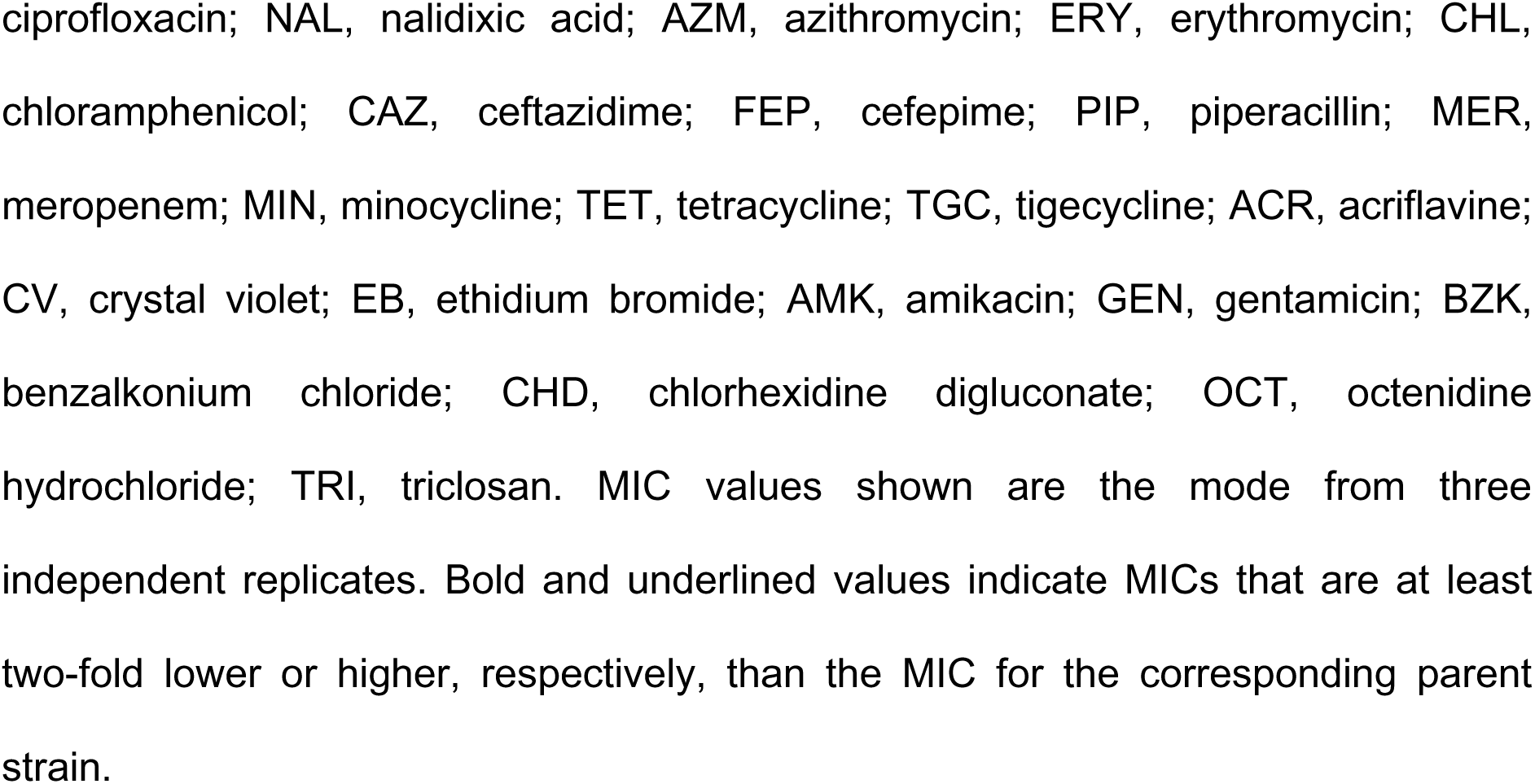
Susceptibility of wild-type *Klebsiella pneumoniae* Ecl8 and ATCC 43816 and their isogenic Δ*ramR* and Δ*acrB* mutant strains to biocides, antibiotics and dyes.

| Antimicrobial <sup>a</sup> | Minimum inhibitory concentration (µg/mL) |  |  |  |  |  |  |
| --- | --- | --- | --- | --- | --- | --- | --- |
| | WT Ecl8 | Ecl8<br>$\Delta ramR$ | Ecl8<br>$\Delta acrB$ | WT<br>ATCC<br>43816 | ATCC<br>43816<br>$\Delta ramR$ | ATCC<br>43816<br>$\Delta acrB$ | ATCC<br>25922 |
| CIP ($\leq 0.25^S$, $> 0.5^R$) | 0.016 | <u>0.125</u> | **0.008** | 0.03 | <u>0.25</u> | **0.008** | 0.008 |
| NAL | 4 | <u>16</u> | **1** | 4 | <u>32</u> | **1** | 2 |
| AZM | 4 | <u>16</u> | **1** | 8 | <u>32</u> | **1** | 4 |
| ERY | 32 | <u>256</u> | **1** | 64 | <u>256</u> | **2** | 32 |
| CHL | 2 | <u>16</u> | **0.5** | 4 | <u>32</u> | **0.5** | 4 |
| CAZ ($\leq 1^S$, $> 4^R$) | 0.06 | <u>0.25</u> | 0.06 | 0.125 | <u>0.5</u> | 0.125 | 0.125 |
| FEP ($\leq 1^S$, $> 4^R$) | 0.016 | <u>0.06</u> | 0.016 | 0.03 | <u>0.125</u> | 0.03 | 0.06 |
| PIP ($\leq 8^S$, $> 8^R$) | 4 | <u>16</u> | 4 | 4 | <u>32</u> | 4 | 1 |
| MER ($\leq 2^S$, $> 8^R$) | 0.03 | 0.016 | 0.03 | 0.03 | 0.06 | 0.03 | 0.03 |
| MIN | 1 | <u>8</u> | **0.125** | 2 | <u>16</u> | **0.125** | 0.5 |
| TET | 1 | <u>4</u> | **0.125** | 1 | <u>8</u> | **0.125** | 0.5 |
| TGC ($\leq 0.5^S$, $> 0.5^R$) | 0.125 | <u>1</u> | **0.03** | 0.125 | <u>2</u> | **0.06** | 0.06 |
| ACR | 32 | <u>128</u> | **4** | 32 | <u>128</u> | **8** | 16 |
| CV | 16 | <u>64</u> | **4** | 16 | <u>64</u> | **4** | 16 |
| EB | 512 | >1024 | **16** | 512 | >1024 | **16** | 512 |
| AMK ($\leq 8^S$, $> 8^R$) | 2 | 2 | 2 | 2 | 2 | 2 | 2 |
| GEN ($\leq 2^S$, $> 2^R$) | 0.5 | 0.5 | 1 | 0.5 | 1 | 1 | 0.5 |
| BZK | 16 | 32 | **2** | 16 | 32 | **4** | 32 |
| CHD | 16 | 32 | **4** | 16 | <u>64</u> | **4** | 0.5 |
| OCT | 2 | 4 | 4 | 4 | 2 | 4 | 1 |
| TRI | 1 | 2 | **0.25** | 1 | 2 | **0.25** | 0.5 |

<sup>a</sup> The clinical breakpoint values from EUCAST are indicated in brackets alongside the antimicrobial, where S and R refer to susceptible and resistant, respectively. CIP,
ciprofloxacin; NAL, nalidixic acid; AZM, azithromycin; ERY, erythromycin; CHL, chloramphenicol; CAZ, ceftazidime; FEP, cefepime; PIP, piperacillin; MER, meropenem; MIN, minocycline; TET, tetracycline; TGC, tigecycline; ACR, acriflavine; CV, crystal violet; EB, ethidium bromide; AMK, amikacin; GEN, gentamicin; BZK, benzalkonium chloride; CHD, chlorhexidine digluconate; OCT, octenidine hydrochloride; TRI, triclosan. MIC values shown are the mode from three independent replicates. Bold and underlined values indicate MICs that are at least two-fold lower or higher, respectively, than the MIC for the corresponding parent strain.

The deletion of *ramR* in Ecl8 and ATCC 43816 reduced susceptibility to a wide range of AcrB substrates compared to their respective wild-type parental strains, indicated by a two-fold or greater increase in MIC values (Table 2). Notably, the MIC values for ceftazidime, cefepime, and piperacillin in the Δ*ramR* strains were two-fold or greater than their respective wild-type strains (Table 2). Tigecycline is one of the last-resort antibiotics for treating *K. pneumoniae* infections. Tigecycline susceptibility in both Ecl8 and ATCC 43816 Δ*ramR* strains was reduced compared to their wild-type strains. According to EUCAST clinical breakpoints, both Ecl8 and ATCC43816 Δ*ramR* strains were clinically resistant to piperacillin and tigecycline (Table 2).

The AcrAB-TolC efflux pump also plays a crucial role in bile acid resistance in Gram-negative bacteria, including clinical *K. pneumoniae* strains (35, 21). Therefore, the growth of the Ecl8 and ATCC 43816 Δ*acrB* strains was monitored in LB broth supplemented with sodium deoxycholate. In 0.5% sodium deoxycholate, the growth of both strains was noticeably reduced, which was reflected in the significantly lower final OD_600_ values in the Δ*acrB* strains compared to their respective parental wild-type strains (Fig. 2A). Compared to Ecl8 Δ*acrB*, the ATCC 43816 Δ*acrB* strain grew better in the presence of 0.5% sodium deoxycholate. In 1% or 2% sodium deoxycholate, the growth of the ATCC 43816 Δ*acrB* strain was further reduced, and both Δ*acrB* strains had a significantly lower final OD_600_ compared to their respective wild-type strains (Fig. 2B and 2C). The Δ*ramR* strains also grew better than their respective wild-type strains in the presence of sodium deoxycholate, indicated by the significantly higher final OD_600_ values (Fig. 2). The growth of the ATCC 43816 Δ*ramR* and Ecl8 Δ*ramR* strains was similar up to 1% sodium deoxycholate, but in the presence of 2% sodium deoxycholate, the ATCC 43816 Δ*ramR* strain grew better than the Ecl8 Δ*ramR* strain, indicated by the higher final OD_600_ value (Fig. 2). This highlights that the AcrAB-TolC efflux pump is crucial for the survival of Ecl8 and ATCC 43816 in sodium deoxycholate.

**Figure 2.**
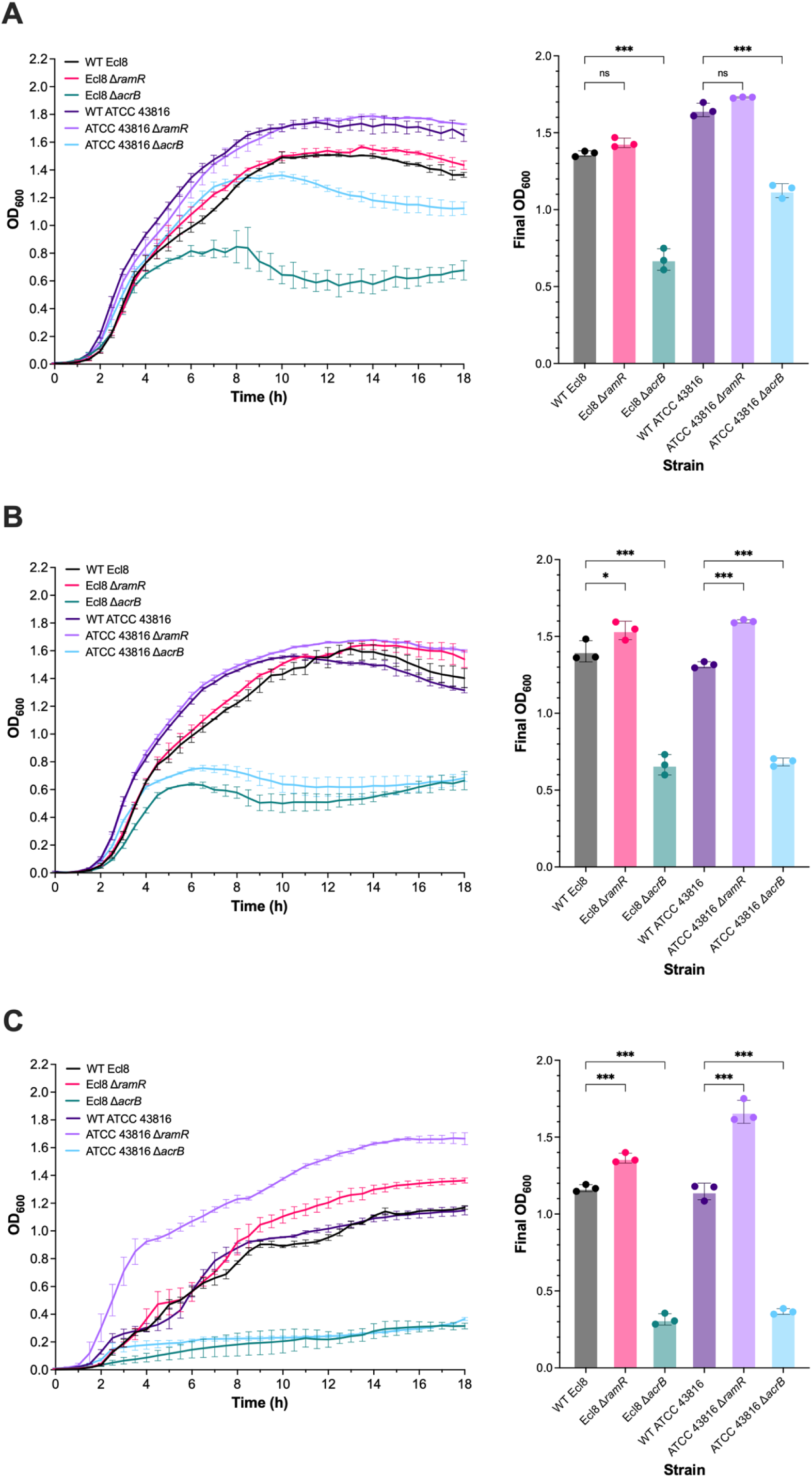
The growth of wild-type *Klebsiella pneumoniae* Ecl8 and ATCC 43816 and their isogenic Δ*ramR* and Δ*acrB* mutant strains in the presence of sodium deoxycholate. The growth kinetics and final OD_600_ values of bacterial cells were measured in LB broth supplemented with **A)** 0.5%, **B)** 1%, and **C)** 2% sodium deoxycholate. The optical density at 600 nm (OD_600_) was measured every 30 min over 18 h at 37 °C with shaking (200 rpm) using a plate reader. The data presented are the mean ± standard deviation of three independent experiments, each consisting of three biological replicates. The final OD_600_ corresponds to the OD_600_ value at the 18 h timepoint. Statistical significance was determined by comparing the wild-type Ecl8 or ATCC 43816 to their isogenic mutant strains using one-way ANOVA, followed by Dunnett’s test to correct for multiple comparisons. Significantly different results are presented and are indicated with * (P ≤ 0.05) or *** (P ≤ 0.001). ns, not significant.

### The effect of *acrB* deletion and overexpression on growth under infection-mimicking conditions in *Klebsiella pneumoniae* Ecl8 and ATCC 43816

As an opportunistic pathogen, *K. pneumoniae* can infect various body sites, including the bloodstream and lungs (36). However, the contribution of the AcrAB-TolC efflux pump to the survival of *K. pneumoniae* under different infection-mimicking conditions has not been investigated. A primary line of defence against invading pathogens is the bactericidal activity of serum (37), and serum resistance enhances pathogenic capacity. Therefore, the survival of the wild-type *K. pneumoniae* Ecl8 and ATCC 43816, as well as their Δ*acrB* and Δ*ramR* mutant strains, in the presence of heat-inactivated or normal human serum over 3 h was measured.

In heat-inactivated serum, there was no difference in survival between wild-type Ecl8 and ATCC 43816 and their isogenic Δ*acrB* and Δ*ramR* mutant strains (Fig. 3A). In normal human serum, the Ecl8 Δ*acrB* strain displayed significantly lower survival compared to its wild-type parent strain, whereas the Ecl8 Δ*ramR* strain showed no significant difference in survival (Fig. 3B). For ATCC 43816, the survival of the Δ*acrB* and Δ*ramR* strains was not significantly different to their parental wild-type strain (Fig. 3B).

**Figure 3.**
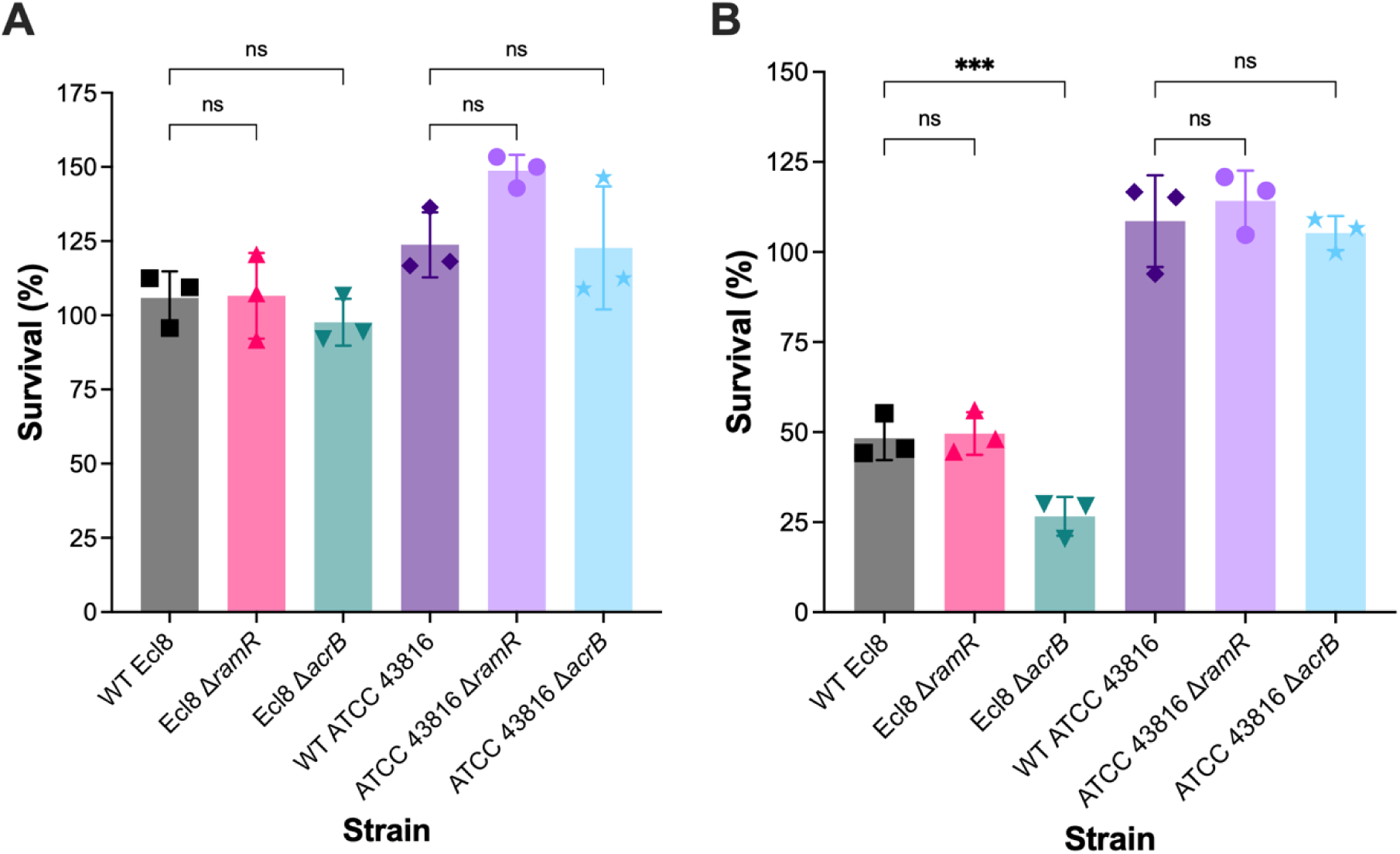
The survival of wild-type *Klebsiella pneumoniae* Ecl8 and ATCC 43816 and their isogenic Δ*ramR* and Δ*acrB* mutant strains in human serum. Survival of wild-type *K. pneumoniae* Ecl8 and ATCC 43816 and their isogenic Δ*ramR* and Δ*acrB* mutant strains in **A)** heat-inactivated serum and **B)** normal human serum. Bacterial cultures were grown to the exponential phase before being added to normal human serum at an inoculum of 1×10^5^ CFU/mL, followed by incubation at 37 °C for 3 h without shaking. Survival was calculated as the number of bacteria treated with heat-inactivated or normal human serum as a percentage of the total number of input bacterial cells. The data presented are the mean ± standard deviation of three biological replicates tested on independent occasions. Statistical significance was determined by comparing the wild-type Ecl8 or ATCC 43816 strains to their isogenic mutant strains using one-way ANOVA, followed by the Holm-Šídák method to correct for multiple comparisons. Significantly different results are presented and are indicated with *** (P ≤ 0.001). ns, not significant.

To cause pneumoniae, *K. pneumoniae* must survive within the lung environment (38). Despite not being a primary cystic fibrosis pathogen, when present, *K. pneumoniae* can be associated with pulmonary exacerbations (39). Therefore, to investigate the contribution of the AcrAB-TolC efflux pump to *K. pneumoniae* growth in both healthy and diseased lung environments, artificial healthy lung media (HLM) and cystic fibrosis lung media (CFLM) were used (32). To recapitulate the lung niche, growth in HLM and CFLM was carried out at 37 °C with 5% CO_2_. As a control, a *P. aeruginosa* strain (PA14) was included as it has been shown to grow in HLM and CFLM (32). As expected, PA14 grew well in HLM and CFLM, indicating stability of the media (Fig. 4). In HLM, Ecl8 and ATCC 43816 Δ*ramR* and Δ*acrB* strains grew at a similar rate to their respective parental wild-type strain, with no significant differences in CFUs at 24 and 48 h time points (Fig. 4A and C). In CFLM, the ATCC 43816 Δ*ramR* and Δ*acrB* strains grew at a similar rate to their wild-type parent strain, with no significant differences in CFUs at 24 and 48 h timepoints (Fig. 4B and D). However, the growth of the Ecl8 Δ*acrB* strain was significantly reduced at 24 and 48 h timepoints compared to wild-type Ecl8, whereas the growth of the Ecl8 Δ*ramR* strain was not significantly different (Fig. 4B and D).

**Figure 4.**
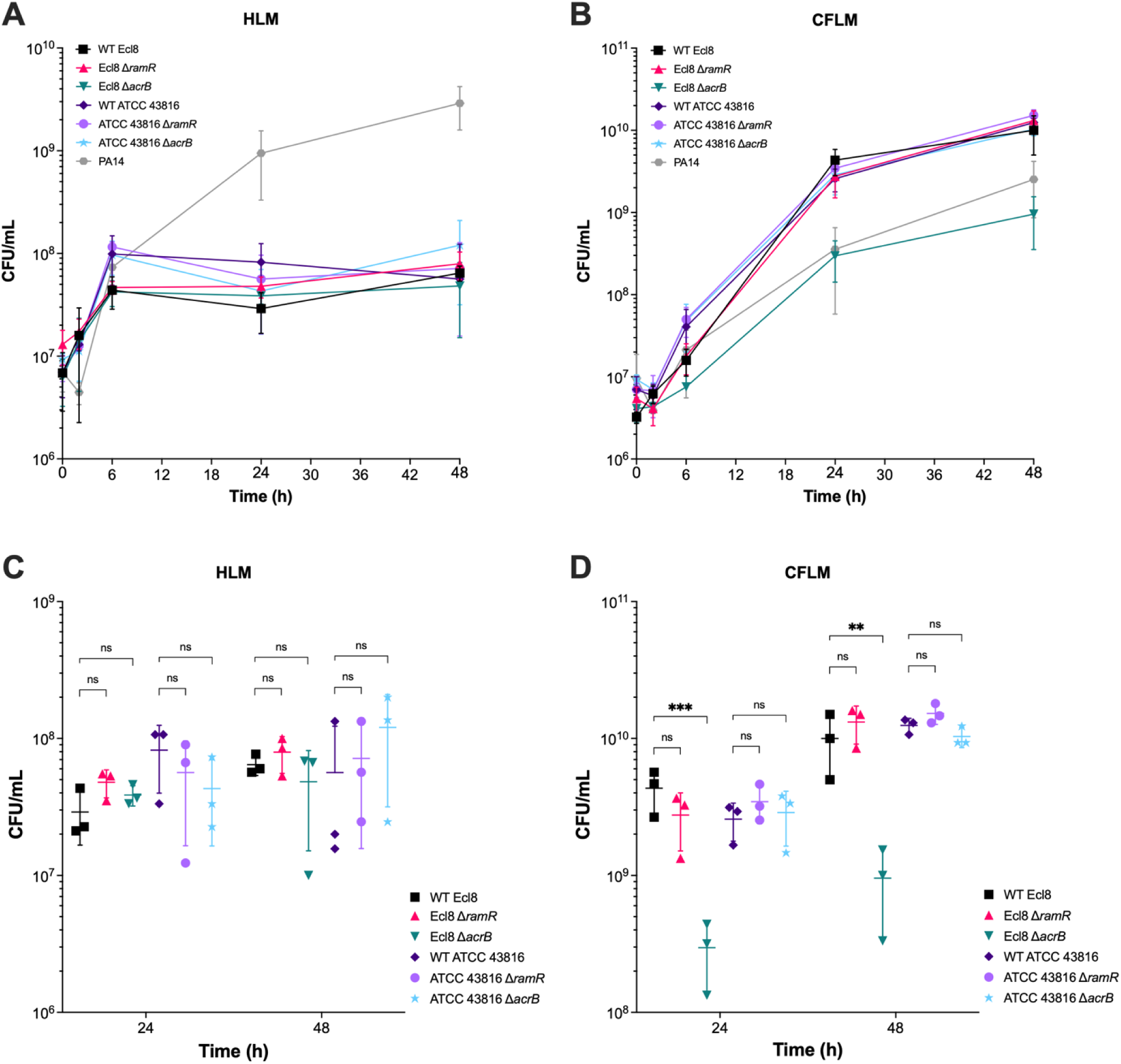
The growth of wild-type *Klebsiella pneumoniae* Ecl8 and ATCC 43816 and their isogenic Δ*ramR* and Δ*acrB* mutant strains in healthy lung media and cystic fibrosis lung media. The growth of wild-type *Klebsiella pneumoniae* Ecl8 and ATCC 43816 and their isogenic Δ*ramR* and Δ*acrB* mutant strains in **A)** healthy-lung media, and **B)** cystic fibrosis lung media. CFU counts were determined by taking samples at 2-, 6-, 24-and 48-hour timepoints. *Pseudomonas aeruginosa* PA14 was included as a control strain. The colony-forming units of the bacterial strains at 24- and 48-hour timepoints in **C)** healthy lung media, and **D)** cystic fibrosis lung media. The data presented are the mean ± standard deviation of three independent experiments, each consisting of three biological replicates. Statistical significance was determined by comparing the wild-type Ecl8 or ATCC 43816 strains at different time-points to their isogenic mutant strains using multiple unpaired lognormal t-tests, followed by the Holm-Šídák method to correct for multiple comparisons. Statistical significance is indicated by ** (P ≤ 0.01), *** (P < 0.001). ns, not significant.

### The effect of *acrB* deletion and overexpression on biofilm formation in *Klebsiella pneumoniae* Ecl8 and ATCC 43816

Biofilm formation in *K. pneumoniae* confers protection against both the host immune response and antibiotics (40, 41). The role of AcrAB-TolC in *K. pneumoniae* biofilm formation has not been explored; therefore, the ability of the *K. pneumoniae* Ecl8 and ATCC 43816 Δ*acrB* and Δ*ramR* mutant strains to form biofilms in tryptic soy broth (TSB), healthy lung media (HLM), and cystic fibrosis lung media (CFLM) was determined using the crystal violet biofilm assay. TSB has been previously shown to support robust biofilm formation by *K. pneumoniae* strains (30). After 72 h in TSB, *K. pneumoniae* Ecl8 formed more biofilm compared to *K. pneumoniae* ATCC 43816 (Fig. 5). The Ecl8 Δ*acrB* strain formed significantly less biofilm, whilst the Ecl8 Δ*ramR* strain was not significantly different compared to wild-type Ecl8 (Fig. 5). Both the ATCC 43816 Δ*acrB* and Δ*ramR* mutant strains formed significantly less biofilm than their parental wild-type strain in TSB (Fig. 5).

**Figure 5.**
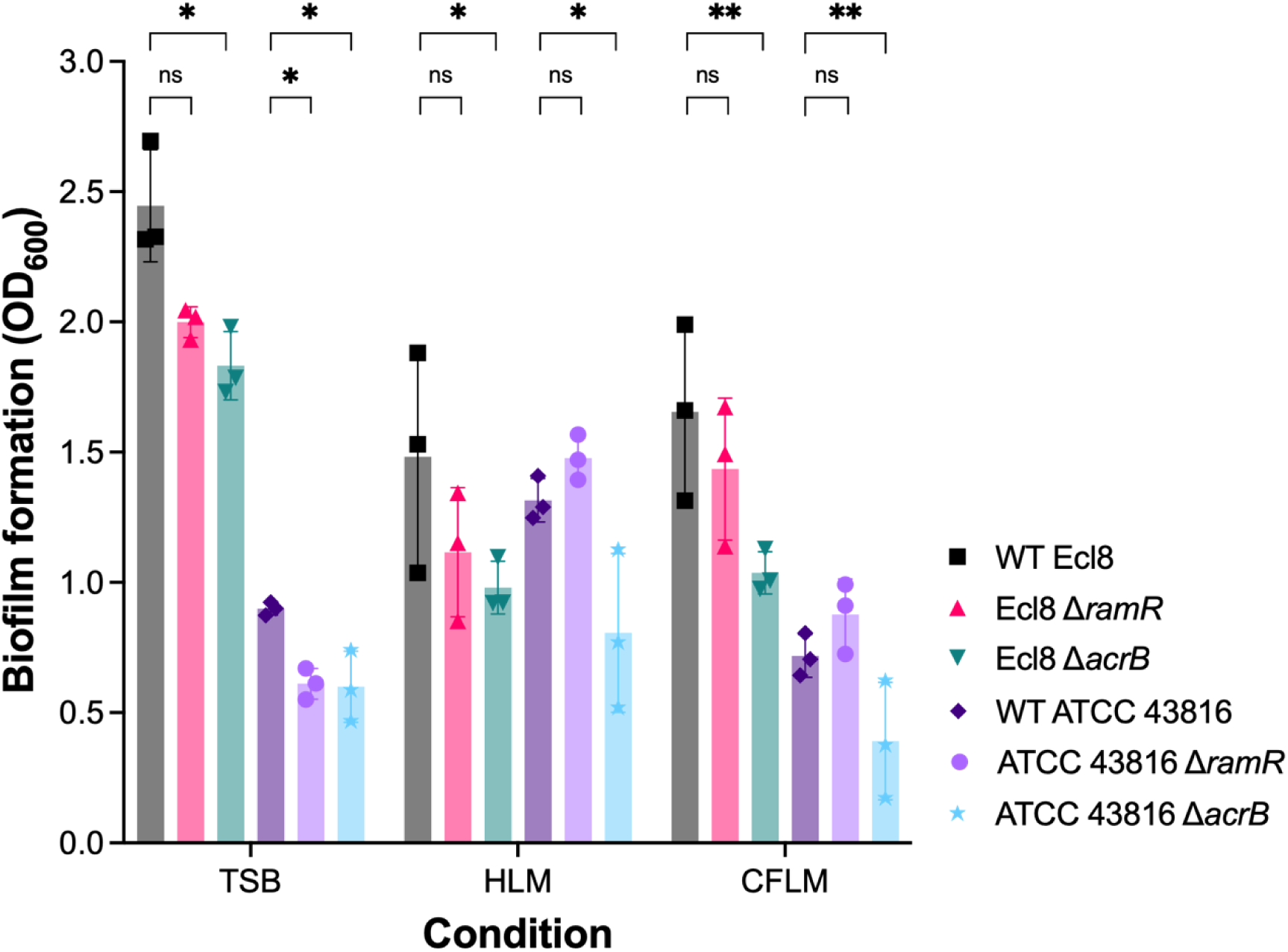
Biofilm formation by wild-type *Klebsiella pneumoniae* Ecl8 and ATCC 43816 and their isogenic Δ*ramR* and Δ*acrB* mutant strains in different media. Biofilm formation (crystal violet staining) at 72 h in tryptic soy broth (TSB), healthy lung media (HLM) or cystic fibrosis lung media (CFLM). The data shown are the mean ± standard deviation of three independent replicates, each consisting of three biological replicates tested in triplicate. Statistical significance was determined by comparing the wild-type strains to their isogenic mutant strains using multiple unpaired t-tests, followed by the Holm-Šídák method to correct for multiple comparisons. Statistical significance is indicated by * (P < 0.05) or ** (P < 0.01). ns, not significant.

In HLM and CFLM, the Ecl8 Δ*acrB strain* formed significantly less biofilm compared to the wild-type Ecl8 strain. In contrast, there was no significant difference in biofilm formation by the Ecl8 Δ*ramR* strain (Fig. 5). However, the reduced biofilm formation in CFLM by the Ecl8 Δ*acrB* strain could be due to its reduced growth (Fig. 4). The ATCC 43816 strain formed more biofilm in HLM and CFLM than in TSB (Fig. 5). Like Ecl8, the ATCC 43816 Δ*acrB* strain formed significantly less biofilm in HLM and CFLM compared to its parental wild-type strain. In contrast, there was no significant difference between the ATCC 43816 Δ*ramR* strain and wild-type (Fig. 5). The reduced biofilm formation by the ATCC 43816 Δ*acrB* strain was likely not due to reduced growth in HLM or CFLM, because it grew similarly to its wild-type parent strain (Fig. 4).

### The effect of *acrB* deletion and overexpression on the virulence of *Klebsiella pneumoniae* Ecl8 and ATCC 43816

In several other Enterobacteriaceae species, the loss of *acrB* has been shown to reduce the ability to invade macrophages and epithelial cells (18, 19, 42). However, the role of the AcrAB-TolC efflux pump on *K. pneumoniae*-host cell interactions has not been investigated. While *K. pneumoniae* is primarily an extracellular pathogen, it has been shown to survive within macrophages and invade lung epithelial cells (43, 44). Therefore, the Ecl8 and ATCC 43816 Δ*acrB* and Δ*ramR* strains were assessed for internalisation and intracellular survival in RAW 264.7 mouse macrophages, as well as for invasion of A549 human lung epithelial cells.

In RAW 264.7 macrophage infection assays, the wild-type Ecl8 and ATCC 43816 strains were internalised after 30 min, with similar internalisation levels at 1 h and 2 h post-infection (Fig. 6A). Intramacrophage survival was assessed by allowing uptake for 30 min, followed by incubation with gentamicin for 24 h. After 24 h post-infection, there was a substantial reduction in the number of recovered wild-type Ecl8 and ATCC 43816 bacterial counts (Fig. 6A), suggesting killing by macrophages. In agreement with a previous study (44), the wild-type strains were not completely killed by macrophages, suggesting intramacrophage survival. Compared to wild-type strains, significantly fewer bacterial counts were recovered for the Ecl8 and ATCC 43816 Δ*acrB* strains after 30 min, 1 h, 2 h, and 24 h post-infection (Fig. 6A). The uptake of the Ecl8 and ATCC 43816 Δ*ramR* strains was not significantly affected after 30 min, 1 h, 2 h, and 24 h compared to their respective wild-type strains (Fig. 6A).

**Figure 6.**
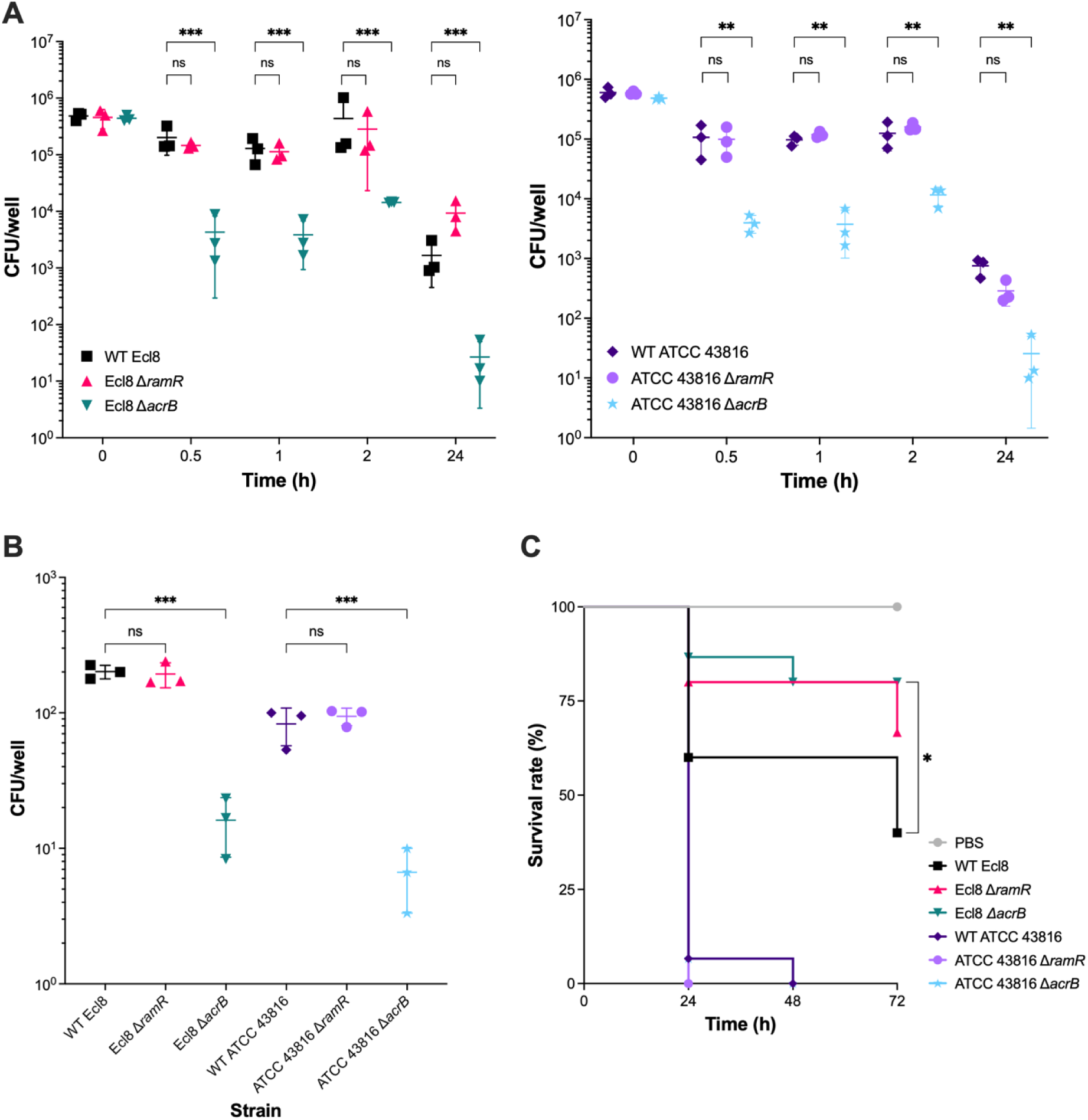
The *in vitro* and *in vivo* virulence of wild-type *Klebsiella pneumoniae* Ecl8 and ATCC 43816 and their isogenic Δ*ramR* and Δ*acrB* strains. **A)** The internalisation and intracellular proliferation of wild-type *K. pneumoniae* Ecl8 and ATCC 43816 and their isogenic Δ*ramR* and Δ*acrB* strains in RAW 264.7 murine macrophages. Macrophage internalisation was determined at 30 min, 1 h and 2 h post-infection. Intramacrophage proliferation was measured 24 h post-infection. Data presented are the mean ± standard deviation of three biological replicates, tested in three technical replicates, on three separate occasions. For each strain, statistical significance was determined by comparing the wild-type strain at different time-points to its isogenic mutant strains using a multiple lognormal t-test, followed by the Holm-Šídák method to correct for multiple comparisons. * (P ≤ 0.05), ** (P ≤ 0.01). **B)** The invasion of A549 human lung epithelial cells by *K. pneumoniae* Ecl8 and ATCC 43816 and their isogenic Δ*ramR* and Δ*acrB* strains after 2 h. Data presented are the mean ± standard deviation of three biological replicates, tested in three technical replicates, on three separate occasions. For each strain, statistical significance was determined by comparing the wild-type strain to its isogenic mutant strains using a one-way ANOVA, followed by Šídák’s multiple comparison testing. *** (P ≤ 0.001), ns, not significant. **C)** Kaplan-Meier survival curves of *Galleria mellonella* over 72 h following inoculation with the *K. pneumoniae* Ecl8 or ATCC 43816 strains. A phosphate-buffered saline injury control is included in grey. For each strain, statistical significance was determined by comparing the wild-type strain to its isogenic mutant strain using the Mantel-Cox test. * (P ≤ 0.05).

Previous work has shown that *K. pneumoniae* 52145R invades A549 lung epithelial cells after 2 h, but the number of viable bacteria steadily decreases over time (43). In agreement, the wild-type Ecl8 and ATCC 43816 strains invaded after 2 h (Fig. 6B). The Ecl8 and ATCC 43816 Δ*acrB* strains were significantly impaired in their ability to invade A549 cells compared to their respective parental wild-type strains (Fig. 6B). The Ecl8 and ATCC 43816 Δ*ramR* strains did not show any significant difference in invasion of A549 cells (Fig. 6B).

Lastly, to assess the role of AcrAB-TolC in in vivo virulence, the *Galleria mellonella* infection model was used. In agreement with previous studies, the wild-type ATCC 43816 strain exhibited a high degree of virulence, resulting in 93% mortality within 24 h and ultimately 100% mortality after 48 h (31). The ATCC 43816 Δ*acrB* and Δ*ramR* strains did not show a significant difference in vivo virulence, with both strains causing 100% mortality by 48 h (Fig. 6C). On the other hand, the wild-type Ecl8 strain was less virulent, causing 60% mortality after 72 h. Compared to wild-type Ecl8, the Δ*acrB* strain was significantly less virulent, whilst the Δ*ramR* strain was not significantly different (Fig. 6C).

## Discussion

Infections caused by MDR *K. pneumoniae* impose a significant global health burden. The archetypal RND efflux pump AcrAB-TolC contributes to MDR, and increasing evidence indicates broader roles in pathogenesis and virulence in Enterobacteriaceae. Therefore, understanding the AMR and virulence determinants in *K. pneumoniae* is crucial for the identification of new drug targets. Here, we identified strain-specific roles for the AcrAB-TolC efflux pump in *K. pneumoniae* growth under infection-relevant conditions and *in vivo* virulence, suggesting that RND efflux pumps do not have the same role within a single species.

The loss of AcrB-mediated efflux activity in *K. pneumoniae* Ecl8 and ATCC 43816 increased susceptibility to a wide range of antimicrobial compounds, consistent with previous studies on other clinical *K. pneumoniae* strains (20, 45, 46). In Enterobacteriaceae, the AcrAB-TolC efflux pump is essential for survival when exposed to bile acids, including sodium deoxycholate (47, 35, 21). Similarly, the loss of AcrB-mediated efflux in Ecl8 and ATCC 43816 also significantly impaired growth when exposed to sodium deoxycholate. Unsurprisingly, this loss did not affect susceptibility to carbapenems or cephalosporins, likely because these antibiotics are poor substrates of the AcrAB-TolC efflux pump (48, 49). However, the known AcrAB-TolC substrate piperacillin (add ref 48) was also not affected by *acrB* deletion, likely due to the presence of the chromosomal *bla*_SHV-1_ gene, which encodes the SHV-1 β-lactamase, found in the vast majority *of K. pneumoniae* strains, including Ecl8 and ATCC 43816 (50). The SHV-1 β-lactamase can hydrolyse penicillins, including piperacillin (51), thereby conferring penicillin resistance regardless of AcrB-mediated efflux (48). Although not used clinically, dyes such as acriflavine, crystal violet, and ethidium bromide are recognised and exported by RND efflux pumps (52). Like *E. coli* and *S*. Typhimurium, the data in this study suggest that the AcrAB-TolC pump in *K. pneumoniae* also exports these dyes.

The overexpression of the AcrAB-TolC pump in Ecl8 and ATCC 43816 Δ*ramR* strains decreased susceptibility to a broad spectrum of antimicrobial agents, consistent with the findings of Majumdar et al. (10). For most antibiotics, except piperacillin and tigecycline, the rise in MIC values was not enough to cause clinical resistance. In the case of piperacillin, resistance in the Δ*ramR* strains was likely due to the increased expression of *acrAB*-*tolC*, working alongside the chromosomally encoded SHV-1 β-lactamase that can hydrolyse penicillins (53). Resistance to tigecycline in clinical *K. pneumoniae* isolates has been linked to the upregulation of *acrB* expression, driven by *ramA* overexpression resulting from inactivating mutations in the *ramR* gene (15, 54). Similarly, tigecycline resistance in the Ecl8 and ATCC 43816 Δ*ramR* strains might be due to increased export of tigecycline by AcrAB-TolC. The elevated AcrAB-TolC activity in the Δ*ramR* strains also resulted in improved growth in the presence of higher concentrations of sodium deoxycholate. However, the loss of *ramR* and the subsequent overexpression of *ramA* also influences the expression of multiple genes, including other MDR-associated genes. (10). Therefore, the phenotype of the Δ*ramR* strains may not be solely due to the overexpression of *acrAB*-*tolC* expression.

The contribution of AcrAB-TolC to growth under infection-related conditions differed between Ecl8 and ATCC 43816. The loss of *acrB* in Ecl8 decreased survival in human serum and growth in cystic fibrosis lung media, whereas in ATCC 43816, deleting *acrB* had no significant effect. Previously, the deletion of *tolC* in ATCC 43816 was found to reduce biofilm formation, capsule production, and serum survival (55). However, the deletion of *tolC* has pleiotropic effects (56); therefore, the phenotypic impact of *tolC* deletion is not necessarily due to the loss of AcrB-mediated efflux. Multiple efflux systems also utilise TolC as an outer membrane channel, meaning the loss of TolC abrogates all TolC-dependent tripartite efflux systems (57). Hence, we interrogated the genomes of Ecl8 and ATCC 43816 using conserved RND protein residues to see whether the ATCC 43816 strain had additional redundant RND or tripartite transporter proteins that could substitute for the loss of AcrB (58). However, both strains had the same number of tripartite RND, MFS and ABC efflux systems, except for KexD, which was only present in Ecl8. Instead, the differences between Ecl8 and ATCC 43816 might be due to variations in capsule expression and production. In *K. pneumoniae*, the capsule contributes to biofilm formation, serum resistance, and growth in nutrient-poor environments (59, 60). ATCC 43816 exhibits the hypermucoviscous phenotype, indicative of high capsule production (61), which likely allows survival in serum and growth in cystic fibrosis lung media in the absence of AcrAB-TolC function. On the other hand, Ecl8 does not exhibit a hypermucoviscous phenotype (Fig. S3), suggesting that it may rely more on AcrAB-TolC as a defence mechanism under stress. The capsule also plays a primary role in defending against *G. mellonella* immunity (62), possibly explaining why the loss of *acrB* was not detrimental to the virulence of ATCC 43816. Previously, the loss of *acrB* in the virulent strain *K. pneumoniae* 52145R was not found to affect capsule polysaccharide or LPS production (20). This suggests that in the virulent strain ATCC 43816, loss of *acrB* also likely had no impact on capsule or LPS production, potentially explaining why it remained virulent in the *G. mellonella* infection model. In both *K. pneumoniae* Ecl8 and ATCC 43816, the loss of *acrB* reduced biofilm formation, consistent with studies in other Enterobacteriaceae species (63, 64). The role of RND efflux pumps in biofilm formation remains unclear. Still, they likely play a multifaceted role, including the export of extracellular polymeric substances and waste metabolites, as well as the dysregulation of biofilm-associated genes (65).

The results of this study suggest that the AcrAB-TolC efflux pump plays an important role in the interaction between *K. pneumoniae* and its host, as well as in strain-dependent differences in virulence. In macrophage infection assays, both wild-type Ecl8 and ATCC 43816 strains were readily internalised, and a fraction of the bacteria persisted within RAW 264.7 cells, which is consistent with earlier reports that macrophages do not completely kill intracellular *K. pneumoniae* (44). In contrast, the Δ*acrB* mutants showed lower levels of uptake, and their survival was markedly reduced after 24 h. This finding suggests that AcrB contributes to bacterial adaptation to the intracellular environment, possibly by preserving cell envelope integrity or reducing susceptibility to macrophage-killing mechanisms. Loss of *ramR*, on the other hand, had little effect on either uptake or survival, suggesting that overexpression of efflux pumps alone does not provide a measurable advantage in this context. A similar picture emerged in epithelial cell invasion assays. Wild-type Ecl8 and ATCC 43816 were able to invade A549 lung epithelial cells, whereas invasion was significantly impaired in their respective Δ*acrB* mutants. The Δ*ramR* strains, however, behaved similarly to their wild-type counterparts, again suggesting that efflux pump overexpression does not enhance host cell entry. The data from the *G. mellonella* infection model highlight that the role of AcrAB-TolC in virulence is not uniform across strains. As expected, the highly virulent ATCC 43816 strain caused near-complete mortality within 48 h, and this outcome was unaffected by deletion of *acrB* or *ramR*. In contrast, the less virulent Ecl8 strain displayed a measurable reduction in pathogenicity when *acrB* was deleted, whereas the loss of *ramR* did not alter the outcome. Taken together, these results suggest a strain-dependent contribution of AcrB, as it appears to contribute to survival and virulence in Ecl8 but is less critical in the more hypervirulent ATCC 43816 background, where other factors may play a more dominant role. A strain-specific role for AcrAB-TolC has also been observed in *Salmonella* Typhimurium virulence. The loss of AcrB function in *S*. Typhimurium DT104 and DT204 did not affect in vivo virulence (66, 67), whereas in *S*. Typhimurium SL1344 and 14028s, loss of AcrB impairs in vivo virulence (17, 18).

In conclusion, our findings suggest that RND efflux pumps can have strain-specific roles within *K. pneumoniae*. Therefore, caution must be taken before assuming that the same efflux pump has the same physiological role within a species.

